# Nature-Inspired Peptide of MtDef4 C-terminus Tail Enables Protein Delivery in Mammalian Cells

**DOI:** 10.1101/2023.09.14.557695

**Authors:** Lucia Adriana Lifshits, Yoav Breuer, Marina Sova, Sumit Gupta, Dar Kadosh, Evgeny Weinberg, Zvi Hayouka, Daniel Z. Bar, Maayan Gal

## Abstract

Cell-penetrating peptides hold great promise as versatile tools for the intracellular delivery of therapeutic agents. Various peptides have originated from natural proteins with antimicrobial activity. In this study, we investigated the mammalian cell-penetrating properties of a 16-residue peptide derived from the C-terminus tail of the *Medicago truncatula* defensin protein, with the sequence GRCRHGFRRRCFCTTHC. We evaluated the ability of this peptide to penetrate multiple types of cells. Our results demonstrate that the peptide efficiently penetrates mammalian cells within minutes and at a sub-micromolar concentration. Moreover, upon N-terminal fusion to the fluorescent protein GFP, the peptide efficiently delivers the GFP into the cells. Despite its remarkable cellular penetration, the peptide has only a minor effect on cellular viability, making it a promising candidate for the development of a cell-penetrating peptide, with potential therapeutic applications.

## Introduction

Cell-penetrating peptides (CPPs) have emerged as powerful tools in the field of biomedical research and therapeutics due to their ability to facilitate the efficient delivery of biomolecules through the cell membranes ^1–3^. The latter selectively regulates the cellular entrance of ions and small molecules and is usually impermeable to larger peptides and proteins. However, CPP amino acid sequences possess unique properties that enable them to overcome the barrier of the cell membrane, allowing for the intracellular delivery of therapeutic molecules, imaging agents, and nucleic acids ^3–5^. Thus, discovering and characterizing new CPPs can significantly advance the development of targeted drug delivery systems, gene therapies, and diagnostic approaches in a broad range of biotechnological fields ^6–11^.

Over 1500 CPPs are currently listed in the CPP2.0 site ^12,13^. Despite their highly divergent sequences and physicochemical properties, these peptides can be broadly divided into cationic, lipophilic, or hydrophobic sequences ^14^. The cellular penetration mechanism of CPPs varies greatly, with cellular internalization predominantly through either endocytosis or direct membrane permeation to the cytosol. Preference for the different pathways relies on the membranal composition, peptide concentration and biophysical characteristics ^15^, all of which directly affect the cellular entry of the CPPs ^16,17^. Additional classification is the source of the peptide sequence, whether originating from a natural protein or synthetically designed ^18,19^. Since the discovery of the transactivating transcription protein from HIV-1 (TAT peptide) ^20^ and the penetration peptide from the Antennapedia homeotic transcription factor ^21^, hundreds of naturally derived CPPs have been characterized. Most of these are cationic peptides that are 6 to 30 amino acids in length ^22–27^. Despite the large number of CPPs, significant drawbacks such as a short lifetime, low penetration efficiency, poor stability, and immunogenicity often hinder the therapeutic and biotechnological applications of CPPs. Thus, further discovery and characterization of novel sequences is of great interest.

Defensins constitute a family of small cysteine-rich antimicrobial peptides of ∼50 amino acids, important in fortifying the plant’s innate immune system by providing defense against various pathogens ^28,29^. Owing to their antimicrobial attributes, plant defensins have attracted substantial interest due to their potential contributions to agriculture and medicine. Defensin antimicrobial characteristics position them as prospective candidates for devising natural alternatives to artificial pesticides. Moreover, their efficacy against human pathogens marks defensins as the basis for the development of a therapeutic alternative in tackling antibiotic-resistant bacteria and fungal infections ^30–32^.

The *Medicago truncatula* defensin 4 (MtDef4) is part of the defensin family that inhibits the growth of plant pathogens ^33^ with a distinct activity profile against various fungal strains ^34,35^. Peptides derived from the C-terminal region of MtDef4 have exhibited potent antifungal properties ^36^. These peptides possess a cationic sequence that enables them to traverse cellular barriers and effectively penetrate fungal cells. The current study explores the cellular penetration of the 16-residue C-terminal peptide of MtDef4 (Def16) into mammalian cells. Our investigation reveals this peptide’s remarkable mammalian cell-penetrating characteristics and its capability to deliver biomolecule cargo into the cells.

## Results

MtDef4 is a 47 amino acid is a defensin protein with potent antifungal properties ^34,35^ (Fig. 1). Its C-terminal tail, characterized by the RGFRRR motif, is pivotal for both its antifungal effectiveness and its ability to penetrate fungal cells. Figure 1 illustrates the structure of MtDef4, highlighting the region containing the 16-residue tail with the sequence GRCRGFRRRCFCTTHC (depicted in dark purple). These specific residues assemble into scorpion toxin-like domain, featuring two cysteine-rich beta strands interconnected by a positively charged loop ^37^.

**Figure 1.**
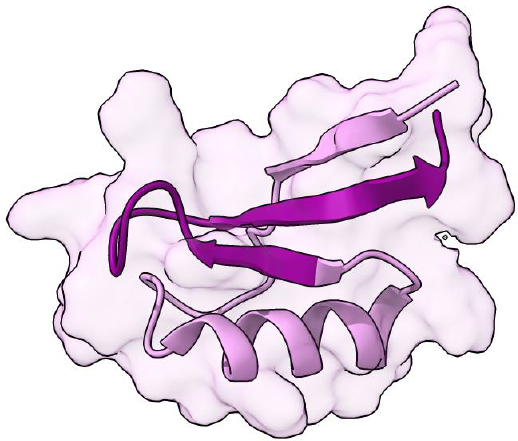
Structure of MtDef4 (PDB: 2LR3) ^33^. The protein’s structure comprises 47 amino acids with a molecular weight of 5.3 kDa. The sixteen residues of the C-terminus tail GRCRGFRRRCFCTTHC (color-coded in dark purple) play a significant role in MtDef4’s function and cellular penetration.

CPPs must show low toxicity to be considered for any biomedical application. Before investigating the ability of the peptide containing the 16-residue C-terminal tail of MtDef4 (Def16) to penetrate mammalian cells and facilitate the delivery of larger proteins, we assessed its impact on cellular viability, establishing a safe and effective concentration range for in-cell applications. To this end, HeLa cells were exposed to varying concentrations of the Def16 peptide for a duration of 24 hours. Cellular viability was evaluated by measuring ATP levels through a luminescence-based assay (Fig. 2A). These data were fitted to a sigmoidal curve, resulting in an IC50 value of 73 µM (Fig. 2B). Thus, only in the tens of micromolar range, the peptide significantly affected cell viability, establishing safe working concentrations for further investigations.

**Figure 2.**
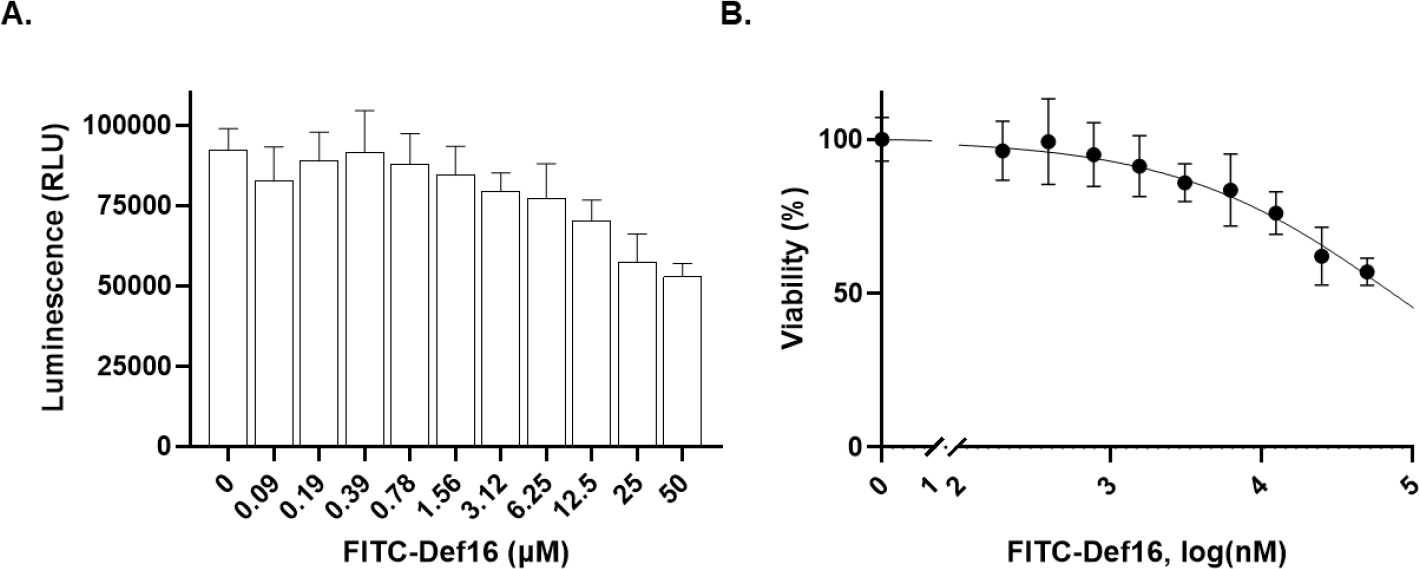
Def16 effect on the cellular viability. HeLa cells were seeded in a 96-well plate at 5,000 cells/well and incubated with variable Def16 concentrations for 24 hours. Cellular viability was determined through the CellTiter assay, quantifying ATP levels. (A) Bar plot showing luminescence values vs. peptide concentrations. (B) Fitting the data into a symmetrical sigmoidal curve yields an IC50 of 73 µM.

To evaluate the capability of Def16 to penetrate mammalian cells, we synthesized the peptide with a fluorescein isothiocyanate (FITC) fluorescent probe at its N-terminus (FITC-Def16). Subsequently, we incubated HeLa cells with varying concentrations of FITC-Def16 for two hours. After thorough washing, we visualized the cells using confocal microscopy imaging. Remarkably, we observed the accumulation and intracellular localization of FITC-Def16 within the cells, even at sub-micromolar concentrations (Fig. 3A,B). FITC fluorescence manifests as green dots distributed within the cells, implying that the peptide gains cellular entry. To test the generality of this peptide, we repeated these experiments with additional cell lines that differ in their membrane composition. Notably, FITC-Def16 also demonstrates similar cell-penetrating capabilities in all other cell types tested, including human gingival fibroblast cells, C2C12 myoblast cells and *A*.*flavus* cells (Fig. S2-S4). To quantify the cellular penetration of FITC-Def16 further, we conducted flow cytometry analysis on HeLa cells following their incubation with varying concentrations of FITC-Def16 (Fig. 3C,D). To ensure that the detected fluorescence signal originated only from peptides internalized within the cells and not bound onto the cells’ membrane, we washed the cells and subjected them to trypsin treatment at 37 °C. This step efficiently eliminates specific and nonspecific extracellular membrane-associated peptides. While the free-FITC control only slightly accumulated in the cell, the FITC-labeled peptide significantly shifted the entire cell population in a concentration-dependent manner. We note that the maximal concentration used was less than 5% of the IC50, a concentration in which we observed no statistically significant effect on cell viability (Fig. 2).

**Figure 3.**
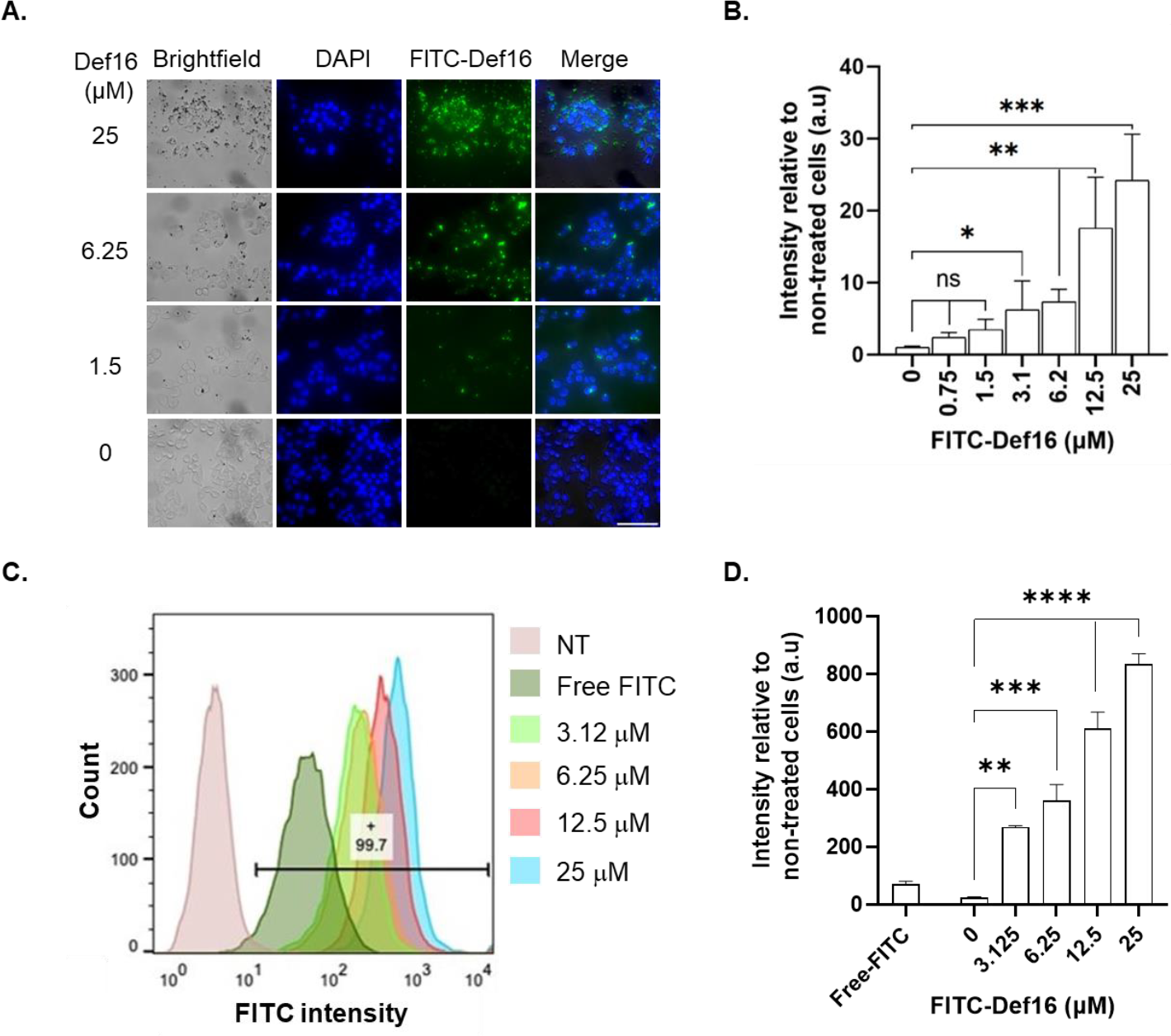
Characterization of Def16 penetration into HeLa cells by confocal imaging and flow cytometry analysis. **(A)** Representative confocal images of HeLa cells treated with variable concentrations of FITC-Def16. Cells were incubated with indicated FITC-Def16 concentrations for two hours at 37 °C, followed by PBS washing and subsequent fixation with 4% PFA and post-fixation staining with DAPI. Images of cells were taken by confocal microscopy, visualizing FITC-Def16 (green) and DAPI (Blue). Scale bar: 100 µm. **(B)** Quantification of FITC-Def16 fluorescence intensity. Bars show the mean green fluorescence intensity relative to non-treated cells and normalized to the number of cells. Values were measured using Color Threshold tab in Fiji. Values shown here are mean ± SD of three replicates. ns -non-significance, *:p<0.05, **:p<0.01, ***:p<0.005. **(C)** Quantification of FITC-Def16 peptide uptake by cells. HeLa cells were incubated with variable FITC-Def16 peptide concentrations for 2 hours, washed, treated with trypsin (0.25%) and subjected to flow cytometry. Diagram representing the uptake of the FITC-containing cells based on the flow cytometry analysis of three independent experiments. Free FITC concentration -25 µM. The horizontal bar and number indicate the percentage of gated FITC-positive cells treated with 25 µM of FITC-Def16. **(D)** Quantitative evaluation of flow cytometry signal intensity. Values were measured using FloJo flow cytometry software. Mean and SD values are calculated from three replicates. **:p<0.005, ***:p<0.0005, ****:p<0.0001.

CPPs that can be uptaken quickly are preferable for biomedical use, as this enables a quicker response time while reducing their dwell time in the extracellular environment, which may modify or degrade them, and reduce the risk of an immunological reaction. To this end, we explored the internalization kinetics of FITC-Def16 into the cells. HeLa cells incubated with a concentration of 6 µM FITC-Def16 were imaged at different time points. A discernible increase in the fluorescence signal, indicative of peptide uptake, was observed during a few minutes with a t_1/2_ of ∼15 minutes (Fig. 4). This places the penetration time of Def16 among other leading CPPs ^38^.

**Figure 4.**
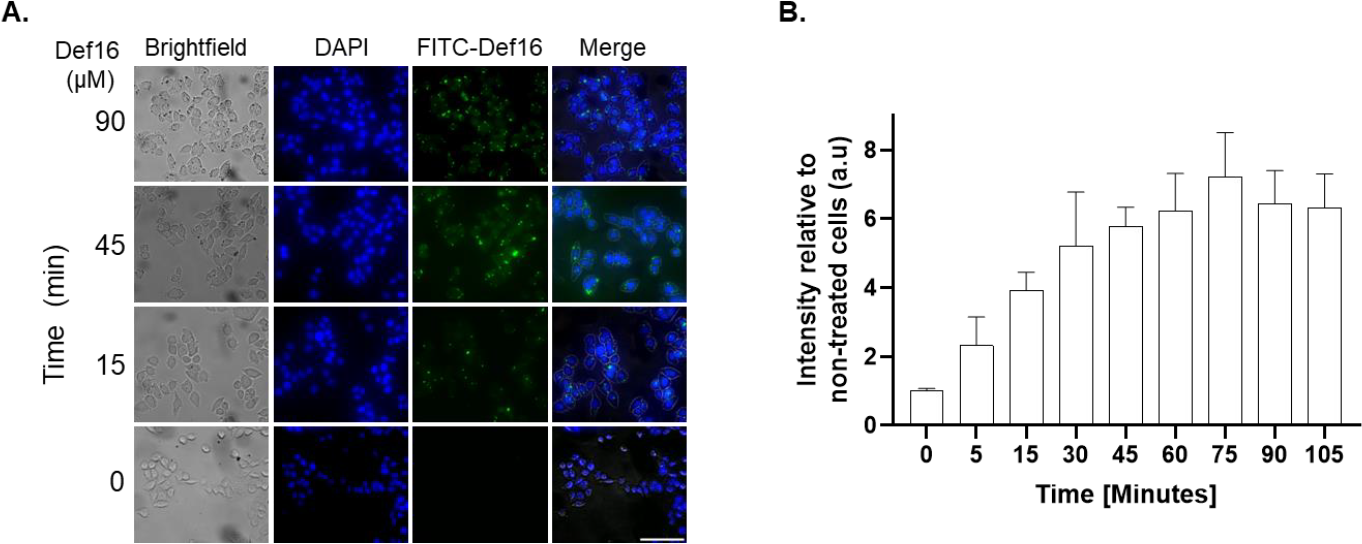
Time response of FITC-Def16 uptake into HeLa cells. **(A)** Cells were cultured with 6 µM of FITC-Def16 at 37 °C, followed by PBS washing and subsequent fixation with 4% PFA and post-fixation staining with DAPI. Images of cells were taken by confocal microscopy visualizing FITC-Def16 (green) and DAPI (Blue). Scale bar: 100 µm. **(B)** Quantification of FITC-Def16 fluorescence intensity. Bars show the mean green fluorescence intensity relative to non-treated cells and normalized to the number of cells. Values were measured using Color Threshold tab in Fiji. Values shown here are mean ± SD of three replicates.

Given the remarkable rate and efficiency of the free peptide cellular penetration, we further investigated its capacity to enhance the delivery of large biomolecules into cells. For that purpose, we evaluated the cellular permeabilization of recombinantly expressed GFP, which was expressed and purified without, and with an N- and C-terminus fusion of Def16 (GFP, Def16-GFP and GFP-Def16, respectively). As expected, GFP alone showed no ability to penetrate cells (Fig. 5A). Surprisingly, while GFP-Def16 minimally penetrated the cells, Def16-GFP penetrated the cells exceptionally well (Fig. 5A,B). This is despite structural predictions indicating that the Def16 is accessible in both proteins, and despite being located not at the very N-terminus, but after a poly-His tag in Def16-GFP (Fig. S1). To gain further insights into the potential cellular mechanism responsible for Def16’s internalization of the cargo protein, we employed staining with a caveolin-1 (Cav-1) antibody. The Def16-GFP protein, shown in the green channel, is only partially co-localized with Cav-1 (red channel), thereby indicating a partial presence of Def16-GFP within endosomes potentially due to endocytosis facilitated through a caveolin-dependent pathway (Fig. 5A, Fig. S3) ^39,40^. Subsequently, we quantified the cellular uptake of the exogenous proteins using flow cytometry analysis. Cells were incubated with each of the recombinant proteins at a concentration of 5 μM for 2 hours. The flow cytometry data further supported our initial observation that Def16-GFP exhibits enhanced cellular uptake compared to GFP-Def16 and GFP alone (Fig. 5C,D, Fig. S5).

**Figure 5.**
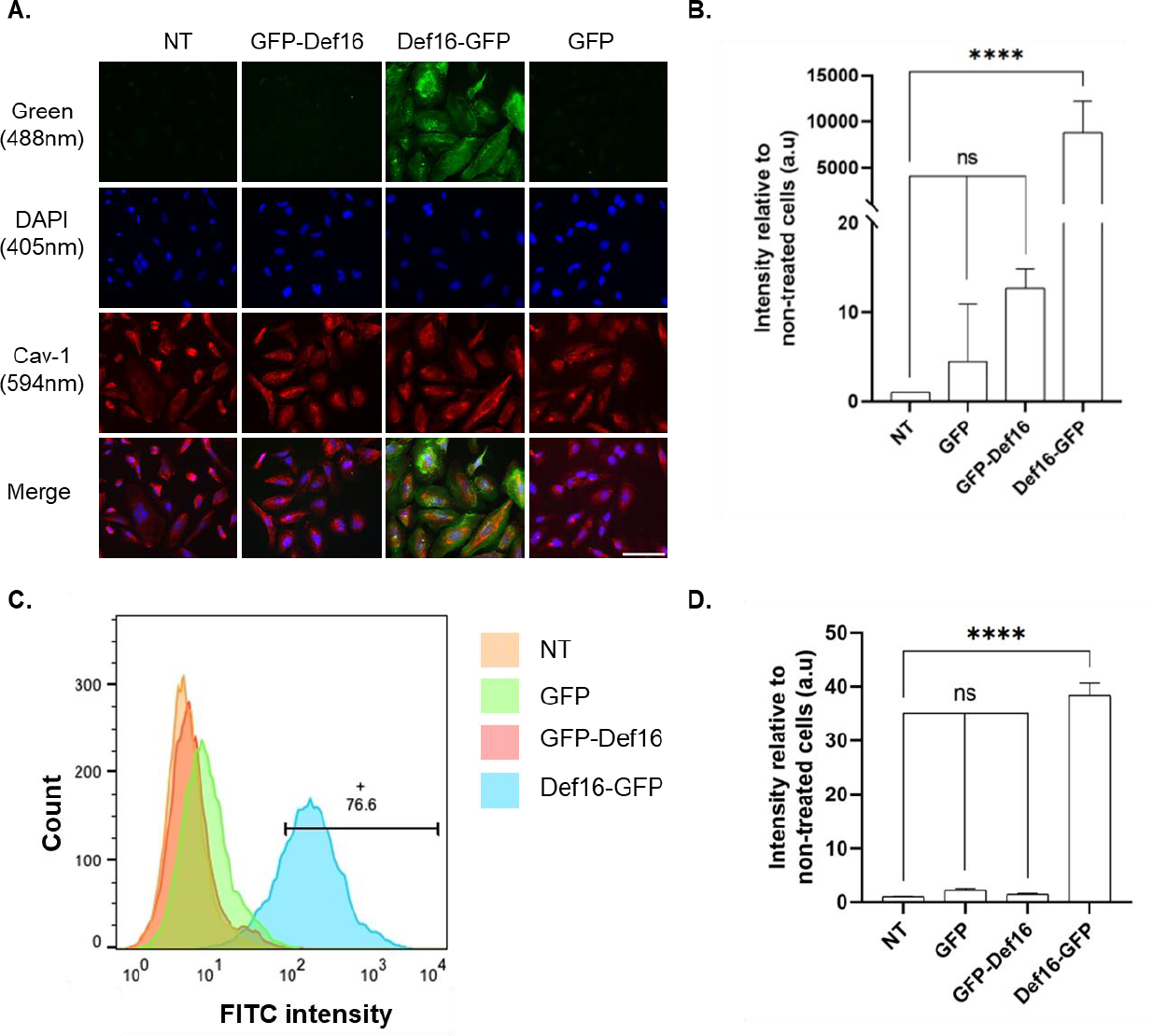
Cellular delivery of GFP conjugated to Def16 peptide. **(A)** HeLa cells were incubated with the recombinant proteins GFP, Def16-GFP and GFP-Def16 at a concentration of 5 μM for 2 hours at **37**°C. Images of cells were taken by confocal microscopy visualizing DAPI (blue), GFP (green) and Caveolin-1 (red) channels. Scale bar: 100 µm. **(B)** Quantification of GFP fluorescence intensity. Bars show the mean fluorescence intensity relative to non-treated cells and normalized to the number of cells. Values were measured using Color Threshold tab in Fiji shown as mean ± SD of three independent replicates. ns -non-significance, ****:p<0.0001. **(C)** Flow cytometry analysis of the cellular uptake of GFP, Def16-GFP and GFP-Def16 to HeLa cells. HeLa cells were incubated with 5 μM of the recombinant proteins for 2h. Diagram representing the uptake of the FITC-containing cells based on the flow cytometry analysis of three independent experiments. NT -Non-treated. The horizontal bar and number indicate the percentage of gated FITC-positive cells. **(D)** Quantitative evaluation of flow cytometry signal intensity. Values were measured using FlowJo flow cytometry software. Mean and SD values are calculated from three replicates.ns -non-significance, ****:p<0.0001.

At low temperatures, membrane lipids undergo a reversible change to a rigid structure, thus limiting energy-dependent endocytosis. Consequently, active endocytosis is anticipated to result in a lower cellular penetration. To delve deeper into the penetration mechanism of Def16 through the cell membrane, we incubated the cells with the proteins at low-temperature conditions (4 °C) and analyzed the data using confocal microscopy. A significant reduction in the penetration of FITC-Def16 and Def16-GFP at 4 °C to HeLa cells is observed (Fig. 6A-B). Flow cytometry further supports these findings, showing a reduced penetration efficiency at 4 °C. Indeed, a reduced percentage of GFP-positive cells was observed at 4 °C (Fig. 6C-D, Fig. S5) in comparison to 37 °C (Fig. 5C, Fig. S5).

**Figure 6.**
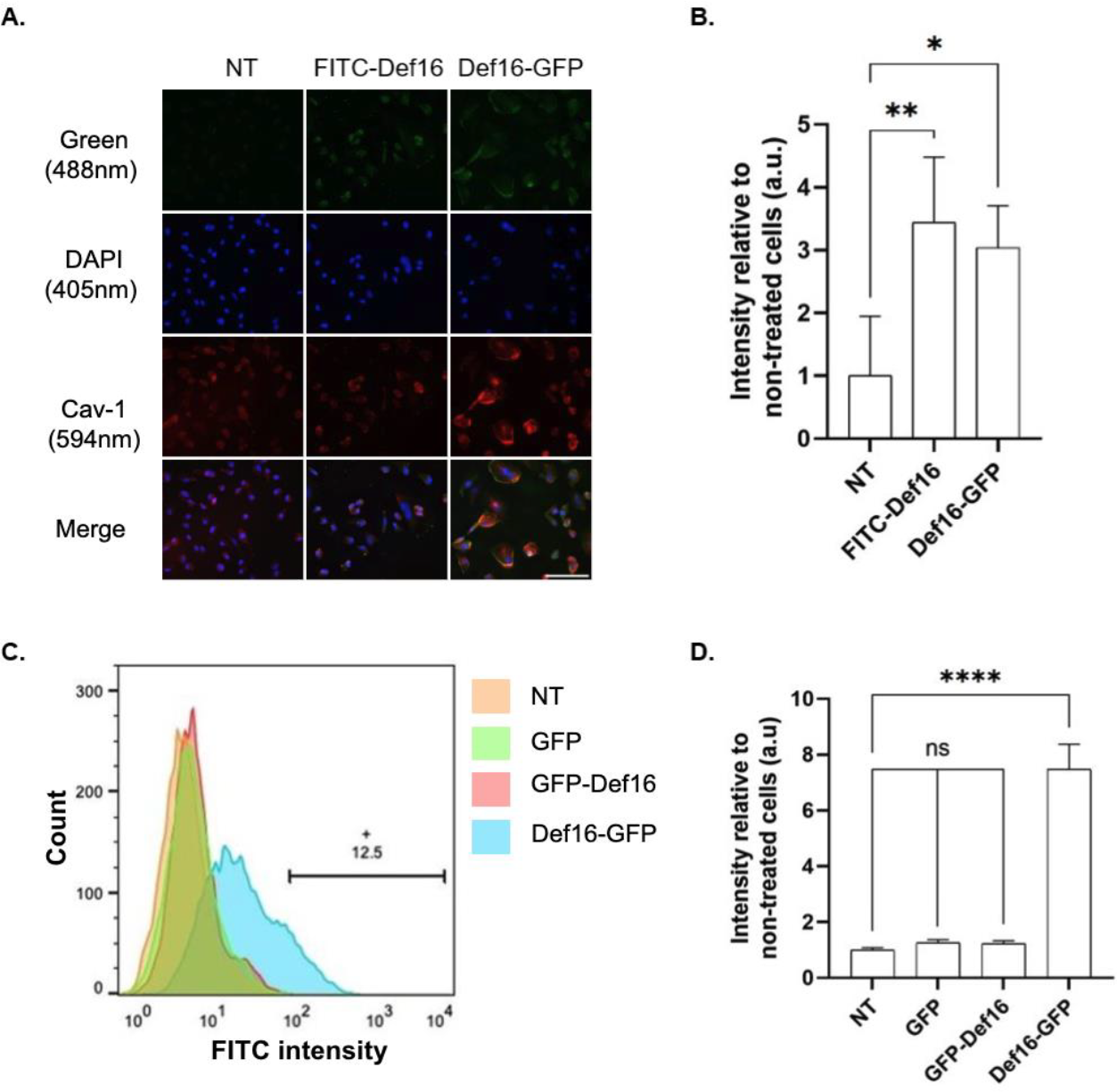
Low-temperature effect on the cellular uptake of Def16 conjugates in HeLa cells. **(A)** Representative confocal images of HeLa cells incubated with 5 μM of FITC-Def16 and Def16-GFP for 2 hours at 4 °C. Images of cells were taken by confocal microscopy, visualizing FITC or GFP (green), DAPI (blue) and Cav-1 (red). Scale bar: 100 µm. **(B)** Quantification of confocal images signal intensity. Bars show the mean green fluorescence intensity relative to non-treated cells and normalized to the number of cells. Values were measured using Color Threshold tab in Fiji. Values show the mean ± SD of three replicates. ns -non-significance, *:p<0.012, **:p<0.0026. **(C)** Flow cytometry analysis of the cellular uptake of FITC-Def16 and Def16-GFP to HeLa cells incubated with 5 μM of the recombinant proteins for 2h at 4°C. Diagram representing the uptake of the FITC or GFP-containing cells based on the flow cytometry analysis of three independent experiments. NT -Non-treated. The horizontal bar and number indicate the percentage of gated GFP-positive cells. **(D)** Quantitative evaluation of flow cytometry signal intensity. Values were measured using FloJo flow cytometry software. Mean and SD values are calculated from three replicates. ns -non-significance, ****:p<0.0001.

## Discussion

CPPs have gained significant attention for their unique ability to facilitate the delivery of various cargoes across cellular membranes. Consequently, the discovery and design of novel CPPs with improved efficiency, specificity, and safety profiles remain highly desirable in biomedical research. Natural antimicrobial peptides, such as defensins, possess inherent cationic membrane-penetrating properties, making them promising candidates for CPP development. This study investigated the cell-penetrating properties of a 16-residue peptide derived from the C-terminus tail of the antifungal defensin protein MtDef4. To characterize Def16’s cellular penetration capabilities, we employed a combination of confocal imaging and flow cytometry analysis, demonstrating its remarkable efficiency in penetrating cells. Our results reveal that this peptide can translocate not only in HeLa cells, but also other cell types such as HGF and C2C12 (Fig. S2, S3). Additionally, we observed penetration into the cell wall of *A*.*flavus* (Fig. S4). which aligns with recent findings in other fungal cells ^36^.

The capacity of CPPs to deliver large biomolecules into mammalian cells is of paramount importance. To assess the effectiveness of Def16 in this regard, we utilized *E. coli* to express a recombinant protein consisting of Def16 fused to the N-terminus of GFP. Remarkably, CPP fusion enabled efficient cellular penetration into cells, despite being ∼14 times larger than Def16 (29.7 kDa including His-tag vs. 2.09 kDa). However, we observed limited cellular transduction when the fusion was made to the C-terminus of GFP (Fig. 5). While the exact mechanism hindering the latter’s cellular entry remains unclear, it may be attributed to structural disparities or the masking of functional segments of Def16, thereby impeding its efficient penetration. Notably, structural predictions by AlphaFold suggest that Def16 remains exposed in both structures. Thus, it could be that excited state population or structural differences that arise only upon contact with the cell membrane drive penetration differences between the two proteins. We also note that the predicted structure of Def16 when fused to GFP (Fig. S1) differs from its resolved structure within MtDef4 protein (Fig. 1). We speculate that the closer spatial proximity of Def16 to the bulk globular structure is essential for an efficient transport across the membrane in Def16-GFP. By contrast, the residual amino acids of the GFP C-terminus, protrude outside the globular structure, distance Def16 from the globular structure, and preclude efficient transport. Consequently, further structural research will be necessary to elucidate the precise orientation of Def16 when fused to a cargo protein and its impact on cellular permeability.

Minimal effect on cell viability is a prerequisite for CPPs considered for most biomedical applications. Our results indicate that efficient cellular penetration of Def16 occurs within a single micromolar concentration range, whereas the cellular viability exhibits an IC50 of 73 µM. This implies that the potential application of the peptide can be at a concentration approximately an order of magnitude lower than its toxic range. This is combined with the fast penetration time, where significant uptake is seen in under 15 minutes and an equilibrium is reached within 30 minutes. Moreover, we quantified nearly 100% permeability into the cells when supplemented with 10 µM of Def16-GFP (Fig. S6). Undoubtedly, conducting additional *in vivo* safety studies to evaluate the peptide’s performance is of high importance. The complexities of circulation and potential degradation may necessitate some alteration, such as higher concentrations to achieve optimal in-vivo penetration, and structural modification to achieve an optimized lifetime. Moreover, CPPs have been associated with potential immunogenic effects, which must be studied further. These studies will not only ensure the safety and biocompatibility of the peptide but also pave the way for its potential translational applications.

While our study primarily focused on the efficacy of cellular penetration, the precise cellular mechanism by which Def16 enters cells remains unclear. Indeed, the determination of an exact cellular uptake mechanism of CPPs is often challenging ^41,42^. In the current study, we observed that the cellular penetration of FITC-Def16 and Def16-GFP is temperature-dependent, with reduced efficiency at lower temperatures. The flow cytometry findings are in qualitative accordance with the observations from confocal microscopy, indicating that the Def16 peptide has the ability to deliver and translocate into diverse cell types and could serve as a possible generalizability as cargo deliver CPP. Moreover, Def16-GFP is partially co-localized with the endosomal marker Cav-1, suggesting cellular entry of the CPP could be driven by endocytosis rather than direct membrane translocation ^43^. However, we can’t rule out the possibility of additional factors preventing Def16 cellular entry at low temperatures, such as the cell membranes becoming less fluid and more rigid. This could hinder the entry of CPPs attempting to cross the membrane under such conditions. In addition, temperature-dependent structural differences of the protein or peptide at high and low temperatures could affect its functionality.

In conclusion, Def16 exhibits significant potential as a cell-penetrating peptide (CPP) with the ability to deliver and internalize biomolecules into various cell types within a relatively short time scale. However, additional aspects such as improved stability against proteases, tissue selectivity, and immunogenicity must be further studied to position it as a promising candidate for biomedical applications, particularly in drug delivery into cells and as a therapeutic peptide.

## Methods

### Peptide synthesis

Def16 peptide with and without FITC was synthesized by Peptide2.0 (Chantilly VA, United States). The peptide was dissolved in DMSO to a stock concentration of 50 mM.

### Cloning of GFP-Def and Def-GFP plasmids

Primers for PCR are listed in Table S1. Cloning of Def16 sequence to pET plasmid containing an N-terminus Hisx6 tag and GFP (Addgene # 29663) was done by whole plasmid amplification using the indicated forward and reverse primers for GFP-Def16 (primers 1 and 2) and Def16-GFP (primers 3 and 4). Following digestion at 37°C with DpnI, the plasmids were phosphorylated by T4 Polynucleotide Kinase and ligated by T4 Ligase (NEB).

### Expression of recombinant proteins

Expression of recombinant proteins (GFP, GFP-Def16 and Def16-GFP) was performed by the transformation of the plasmid into *E. coli* BL21 (DE3). Bacterial growth was carried out at 37 °C and 200 rpm until OD_600_ was ∼0.8, and protein expression was induced by supplementing 1 mM Isopropyl β-d-1-thiogalactopyranoside (IPTG) for 16 hours at 25°C. Following three sonication cycles of three minutes each, the bacterial lysate was centrifuged at 16,000xg for 30 min. The supernatant was passed onto a nickel column, and after extensive wash, the protein was eluted with 300 mM imidazole. Following buffer exchange to phosphate-buffered saline (PBS), the protein was kept frozen with the addition of 50% glycerol. Protein concentrations were assessed using NanoDrop (NanoDrop OneC).

### Cell cultures

Experiments with primary human gingival fibroblasts (hGFs) were approved by the Tel Aviv University institutional review board (IEC No. 0001006-1). Informed consent was obtained from all patients. Human gingival fibroblasts (HGF) derived from masticatory mucosa were cultured as previously described ^44,45^. Briefly, HGF cells were thawed with the appropriate medium and the third passage was examined. Cells were grown on a Dulbecco’s modified Eagle’s medium (DMEM) with 10% fetal calf serum, 100 U/mL penicillin, and 100 mg/mL streptomycin, 1% MEM-Eagle, Hanks’ salts and 1% Sodium Pyruvate. The cervical cancer cell line HeLa and C2C12 myoblast cells were cultured in a standard culture medium (Dulbecco’s modified Eagle’s medium (DMEM)) with 10% fetal calf serum, 2 mM glutamine, 100 U/mL penicillin, and 100 µg/mL streptomycin.

### Immunofluorescence confocal microscopy

For microscopy images, HeLa and C2C12 cells were cultured the day prior to the experiment on round-glass coverslips in a 24-well plate or in a 96-well glass bottom plate and incubated with medium supplemented with the indicated concentrations of peptide/protein at 37°C under 5% CO2. Cells were then washed three times with cold PBS, followed by a cleansing process utilizing light shaking together with three washes of PBS, then three washes with 25mM of Hepes to eliminate nonspecific protein binding. The cells were fixed with 4% paraformaldehyde (PFA) in PBS for 20 minutes and washed three times with PBS to remove the PFA. All samples were stained with 1 mg/mL 4′,6-diamidino-2-phenylindole (DAPI Staining Solution ab228549, Abcam). Cav-1 (ab2910, Abcam), β-tubulin (05-661, Merck). Images were taken directly from the 96-well plate glass bottom, while coverslips from the 24-well plate were transferred into a glass slide and placed on the plate with sodium phosphate buffer mounting medium (25% Mowiol, 50% glycerol, 50% sodium base pH 8.0). Images were taken using a ZEISS LSM 900 in confocal mode, utilizing Zen 3.1 software. Fluorescence readouts were obtained for blue, green, bright red and far-red at ex/em of 358/461nm, 495/519 nm, 561/594nm and 652/668nm, respectively.

### Image analysis

Image acquisition and analysis were carried out with Fiji/ImageJ (version 2.14.0/1.54f), https://imagej.net/software/fiji/downloads. The fluorescence intensity of FITC labeled peptide/recombinant protein was quantified using the Color threshold option, which monitored the fluorescence values and normalized by cell number.

### Cellular viability

HeLa cells were seeded the day prior to incubating them with the peptides. Subsequently, cells were incubated with the indicated concentrations of the labeled peptide for 24 and 48 hours. Cellular ATP levels were quantified using the CellTiter-Glo® assay (Promega, G7570). The reagent was mixed at a 1:1 ratio with supernatant from the treatment plate. The mix was incubated for 10 min at room temperature on the shaker, followed by luminescence measurement using a plate reader (Biotek Synergy H1).

### Structural modeling and visualization

GFP, Def16-GFP, GFP-Def16 protein structure predicted by AlphaFold2 with MMseqs2 (https://colab.research.google.com/github/sokrypton/ColabFold/blob/main/AlphaFold2.ipynb). Structural visualization of GFP proteins and *Medicago truncatula* (PDB ID: 2LR3) was performed using UCSF ChimeraX ^46^.

### Flow cytometry

Cells were seeded at a density of 3×10^5^ cells per well in a 6-well plate and grown at 37 °C in a 5% CO2 atmosphere in DMEM supplemented with 10% (v/v) FBS for 18 h. Cells were then incubated with indicated concentrations of peptide/proteins in fresh DMEM at 37°C or 4°C for 2 h, followed by washing three times with PBS. Cells were treated with Trypsin (100 μl/well of 0.25 %Trypsin). 1 ml PBS was added to the wells with the trypsinized cells, centrifuged, and the supernatant was discarded, and the cells were resuspended in cold PBS with 2.5% FBS and analyzed by flow cytometry. FITC/GFP signal was monitored using Flow cytometer S1000EXi (Stratedigm, San Jose, CA). 10,000 cells were analyzed per single read. Dead cells were gated out from the analysis. Data were analyzed by CellCapTure and FlowJo v10.

### Statistical analysis

Data analysis, bar plots and statistical evaluations were performed using GraphPad Prism version 9.0.0 (www.graphpad.com). One-way ANOVA followed by Dunnett’s multiple comparisons post hoc test.

## Acknowledgments

This research was supported by the Gertner Institute for Medical Nano System at Tel Aviv University, the Israeli Science Foundation (Daniel Z Bar, ISF 632/20) and Bountica Ltd funded research for M.G and Z.H. L.A.L is grateful for Ph.D. scholarship financial support from the ADAMA Center in TAU. pET His6 GFP TEV LIC cloning vector (1GFP) was a gift from Scott Gradia (Addgene plasmid # 29663).

## Data availability statement

All the data generated and analyzed during the current study is available in the manuscript.

## Author Contributions

L.A.L, M.S, and Y.B executed the experiments and analyzed the data. D.K executed the experiments on HGF cells. S.G expressed and purified recombinant proteins. L.A.L analyzed the data and wrote the manuscript. Z.H, E.Y and D.Z.B assisted in manuscript editing, experimental planning and microscopy imaging. M.G wrote the manuscript, conceived the original research and supervised the research. All authors gave their final approval and agreed to be accountable for all aspects of the work.

