## Supplementary material for "Nature-Inspired Peptide of MtDef4 C-terminus Tail Enables Protein Delivery in Mammalian Cells": Fig. S

**Supplementary Table S1**

| Primer name | Sequence | Plasmid |
|---|---|---|
| Primer 1 | F: 5'-CGTCGCCGCTGCTTCTGCACCACGCATTGTTAACGGATCCGAATTCGAG-3' | GFP-Def |
| Primer 2 | R: 5'-AAAACCACGGCAACGGCCACTGCCTCCCCCCTTGTACAGCTCGTCCATGCCGAG-3' | GFP-Def |
| Primer 3 | F: 5'-CGCTGCTTCTGCACCACGCATTGTGGGGGAGGCAGTGTGAGCAAGGGCGAGGAGCT-3' | Def-GFP |
| Primer 4 | R: 5'-GCGACGAAAACCACGGCAACGGCCAGAACCATGGTGATGGTGATGGTGAGAAG-3' | Def-GFP |

**Supplementary Table S2**

| Protein name | Sequence |
|---|---|
| GFP | MGSSHHHHHHGSSVSKGEELFTGVVPILVELDGDVNGHKFSVRGEGEGD<br>ATNGKLTLKFICTTGKLPVPWPTLVTTLTYGVQCFSRYPDHMKQHDFFKSA<br>MPEGYVQERTISFKDDGTYKTRAEVKFEGDTLVNRIELKGIDFKEDGNILGH<br>KLEYNFNSHNVYITADKQKNGIKANFKIRHNVEDGSVQLADHYQQNTPIGD<br>GPVLLPDNHYLSTQSKLSKDPNEKRDHMVLLEFVTAAGITLGMDELYKGIE<br>ENLYFQSNIGSG |
| Def16-GFP | MGSSHHHHHHGSGRCRGFRRRCFCTTHCGGGSVSKGEELFTGVVPILVE<br>LDGDVNGHKFSVRGEGEGDATNGKLTLKFICTTGKLPVPWPTLVTTLTYGV<br>QCFSRYPDHMKQHDFFKSAMPEGYVQERTISFKDDGTYKTRAEVKFEGDT<br>LVNRIELKGIDFKEDGNILGHKLEYNFNSHNVYITADKQKNGIKANFKIRHNV<br>EDGSVQLADHYQQNTPIGDGPVLLPDNHYLSTQSKLSKDPNEKRDHMVLL<br>EFVTAAGITLGMDELYKGIEENLYFQSNIGSG |
| GFP-Def16 | MGSSHHHHHHGSSVSKGEELFTGVVPILVELDGDVNGHKFSVRGEGEGD<br>ATNGKLTLKFICTTGKLPVPWPTLVTTLTYGVQCFSRYPDHMKQHDFFKSA<br>MPEGYVQERTISFKDDGTYKTRAEVKFEGDTLVNRIELKGIDFKEDGNILGH<br>KLEYNFNSHNVYITADKQKNGIKANFKIRHNVEDGSVQLADHYQQNTPIGD<br>GPVLLPDNHYLSTQSKLSKDPNEKRDHMVLLEFVTAAGITLGMDELYKGGG<br>SGRCRGFRRRCFCTTHC |

**Supplementary Figure S1**

Recombinant GFP with Def16 at either the N- or C-terminus (Def16-GFP or GFP-Def16, respectively) was cloned into the pET28 vector and expressed in *E. coli* BL21 (DE3) cells. All plasmids feature an N-terminus Hisx6 tag. Figure S1-A provides an illustration of the expressed constructs. Figure S1-B illustrates the structural model of the proteins. Detailed protein sequences are found in Table S2.

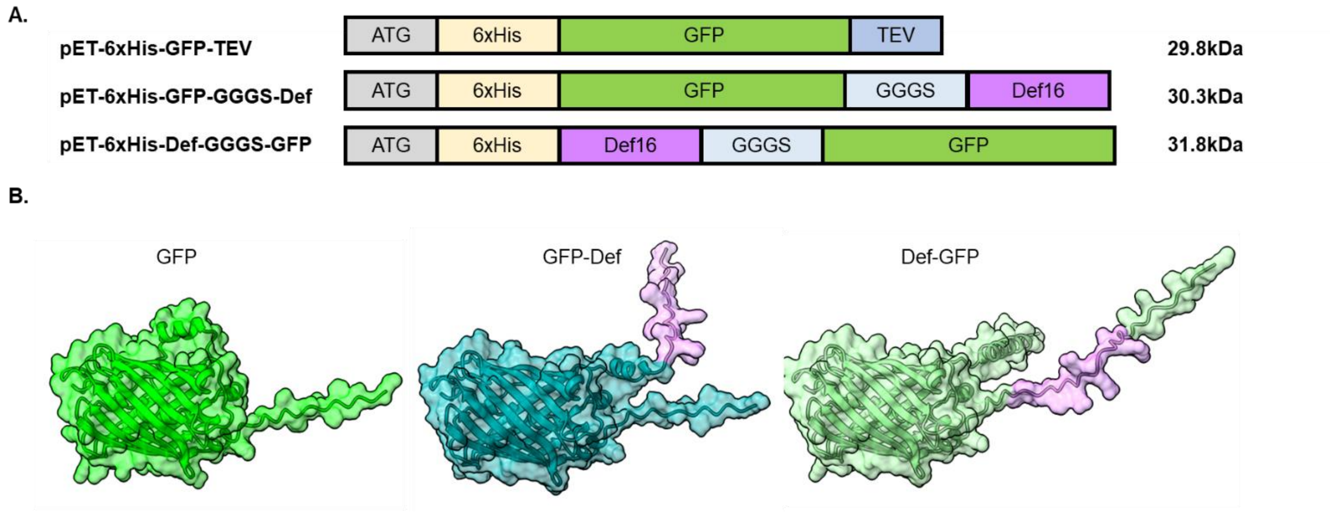


**Figure S1. Expression of recombinant GFP protein with N- and C-terminus Def16 tag. (A)** Illustration of protein constructs. **(B)** Comparison of AlphaFold models of GFP-tagged proteins with the Def16 peptide. Surface structure representations created with ChimeraX software.

**Supplementary Figure S2**

To assess Def16's capacity for cell penetration across different cell types, we subjected HGF cells to various concentrations of FITC-Def16. After two hours of incubation and thorough washing, we examined the cells using confocal microscopy imaging. Figure S2 presents a series of images, including representative brightfield, DAPI (blue), FITC (green), and merged images. The images show the accumulation of FITC-Def16 within the cells.

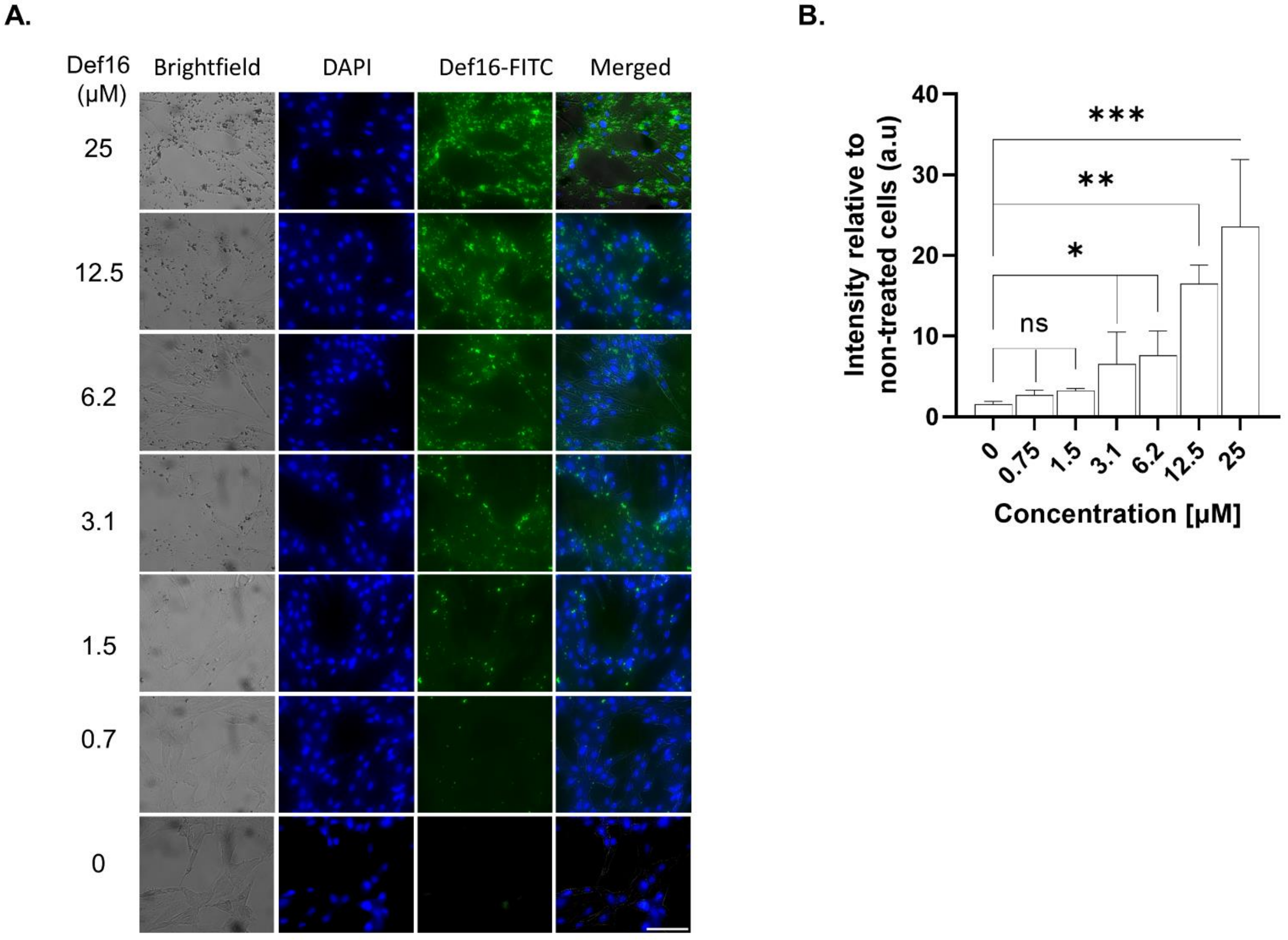


**Figure S2. FITC-Def16 Penetration into HGF Cells. (A)** HGF cells were cultured for two hours with varying concentrations of FITC-Def16 and subsequently stained with DAPI. Confocal microscopy images were captured to visualize FITC-Def16 (green) and DAPI (blue) in cells attached to a slide. Scale bar: 100 μm. **(B)** Mean green fluorescence intensity relative to non-treated cells and normalized to the cell count. The signal was quantified using the Color Threshold tool in Fiji. Significance levels are denoted as follows: *p<0.05, **p<0.01, ***p<0.005.

**Supplementary Figure S3**

To investigate the intracellular protein uptake facilitated by the Def16 peptide across various cell lines, we treated C2C12 myoblasts with either 5 μM of FITC-Def16 peptide or GFP-tagged proteins for a duration of 2 hours. Subsequently, the cells underwent three PBS washes, followed by fixation using 4% PFA. Finally, they were mounted on a glass slide for evaluation through confocal microscopy. Consistent with the results presented in Figure 3, the fluorescence microscopy images herein provide further confirmation of the efficient cellular uptake of FITC-Def16. Additionally, these images reveal partial colocalization with the endosomal marker, Cav-1. A similar outcome was observed in HeLa cells (Figs. 5 and 6), where Def16 effectively demonstrated its competence in internalizing GFP cargo into the cells.

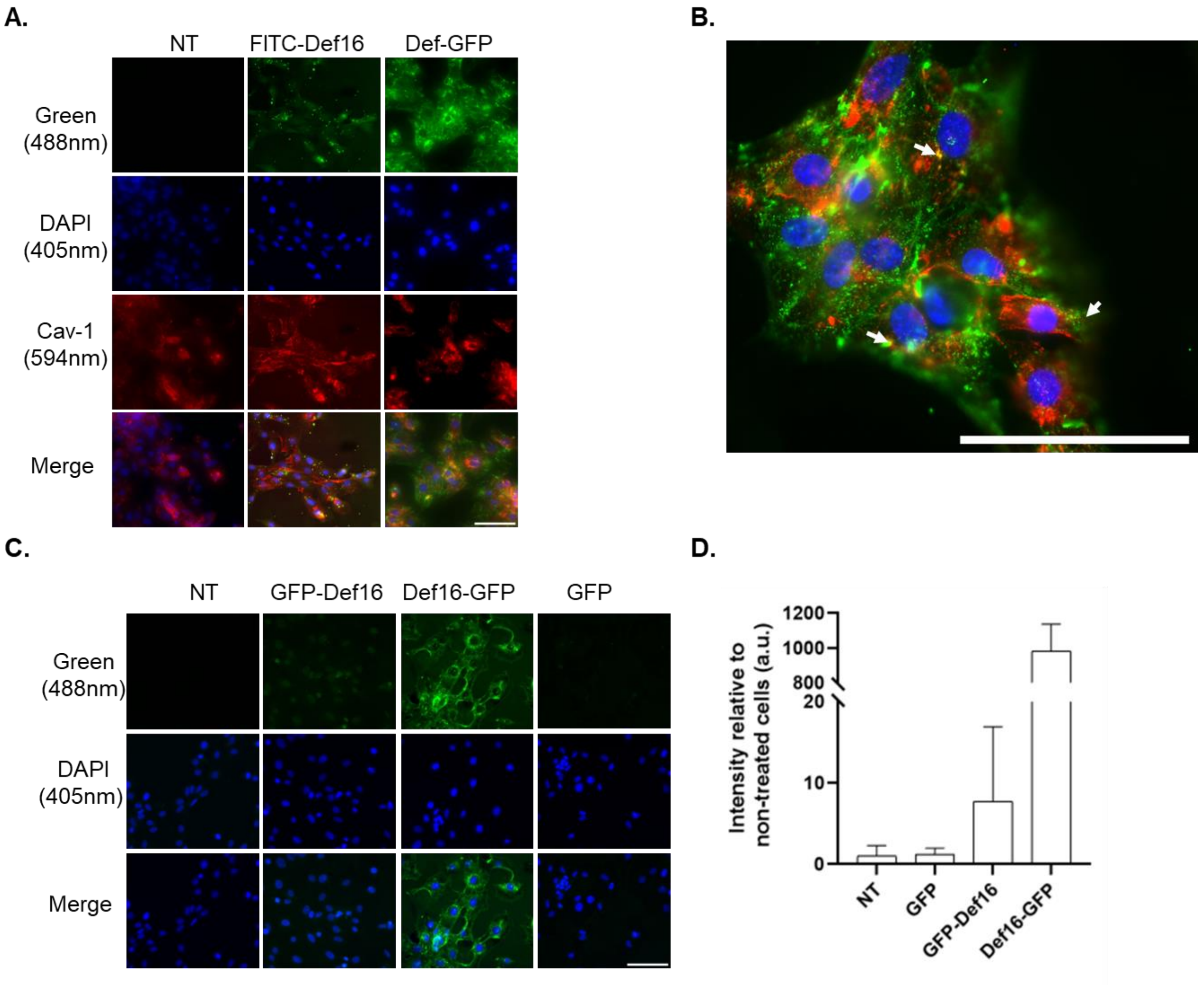


**Figure S3. Facilitation of GFP Protein Delivery by Def16 in C2C12 Cells. (A)** Partial colocalization of FITC-Def16 and Def16-GFP in C2C12 cells with endosomal marker. Cells were incubated with 5 μM of FITC-Def16 or Def16-GFP for two hours, fixed, mounted, and scanned by a confocal microscope. The images visualize FITC or GFP (green), DAPI (blue), and Cav-1 (red). Scale bar: 100 µm. Colocalization is indicated when similarly shaped structures appear yellow in the Merge. **(B)** Representative figure of Def16-GFP colocalized with Cav-1 (white arrowheads), taken with a 63x Objective Lens. Scale bar: 100 µm. **(C)** Confocal microscopy images were captured to visualize GFP tagged with the Def16 peptide (green) and DAPI (blue) in cells attached to a slide. Scale bar: 100 µm. **(D)** Mean green fluorescence intensity relative to non-treated cells and normalized to the cell count is presented in this section. Measurement values were obtained using the Color Threshold tool in Fiji.

**Supplementary Figure S4**

To assess the cellular uptake of FITC-Def16, we incubated fungal cells, *A.flavus* with varying concentrations of FITC-Def16. Figure S4 displays representative images, including brightfield, DAPI (nuclear DNA stain), green fluorescence (FITC), and merged images. Notably, these images clearly reveal the accumulation of Def16-GFP within the cells.

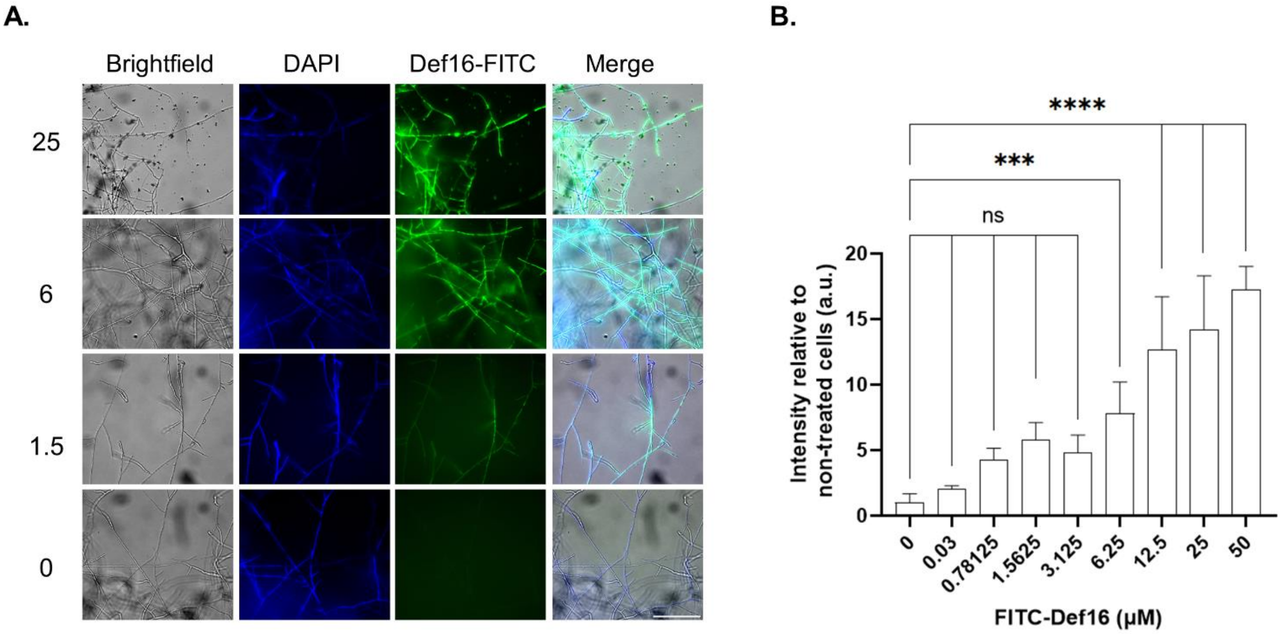


**Figure S4. FITC-Def16 penetrates into *A.flavus*. (A)** Fungal cells were cultured for two hours with variable concentrations of FITC-Def16 and stained with DAPI. Images of cells attached to a glass slide were taken by confocal microscopy visualizing FITC-Def16 (green) and DAPI (Blue). Scale bar: 100µm. **(B)** Mean green fluorescence intensity relative to non-treated cells and normalized to the number of cells. Values were measured using Color Threshold tab in Fiji. , ***:p<0.005, ****:p<0.0001.

### *A. flavus* culture and imaging

*A. flavus* A11 strain was generously provided by the laboratory of Prof. Nir Ashrov (TAU). Spores were initially seeded on YAG-agarose 10 cm plates and allowed to grow for 3-4 days in a humidified incubator at 30 °C. After sporulation, fresh spores were collected using 5 ml of double-distilled water (DDW) containing 0.2% Tween. The spores were then centrifuged at room temperature for 5 minutes at 4,000 RPM and suspended in 5 ml of DDW to create the *A. flavus* fresh stock. To determine the spore concentration in the fresh stock, a 1,000-fold dilution was made with DDW, and the spores were counted using a cytometer. For microscopy purposes, 100 µL of fungal spores were cultured in 10 mL of fungal media (consisting of RPMI-1640 Medium, 100 U/mL penicillin, 100 mg/mL streptomycin, and 165 mM MOPS) in a 10 cm round dish for overnight hyphal growth at 30 °C. The following day, hyphae were collected, resuspended, and cultured with 1 mL of fungal media on top of round-glass coverslips placed in a 24-well plate. Fungi were then incubated for 16 hours at 30 °C. Subsequently, the fungal hyphae were treated with the indicated concentrations of FITC-Def16 peptide and incubated for 2 hours at 30°C. The treated fungus was washed twice with cold PBS, followed by gentle centrifugation at 4,000 rpm for 10 minutes. Finally, the fungus was fixed with 4% paraformaldehyde (PFA), washed with PBS, and examined using confocal microscopy imaging.

**Supplementary Figure S5**

To explore a potential mechanism for the cellular entry of Def16-GFP and GFP-Def16, we conducted an experiment in which HeLa cells were incubated with the proteins for two hours at 37 ℃ and 4 ℃. Our aim was to assess the extent of permeabilization of the proteins and investigate the impact of temperature on cellular permeability. To quantify the levels of permeabilized proteins, we performed a flow cytometry analysis on HeLa cells exposed to varying temperature treatments, enabling us to compare their effects on cellular permeability.

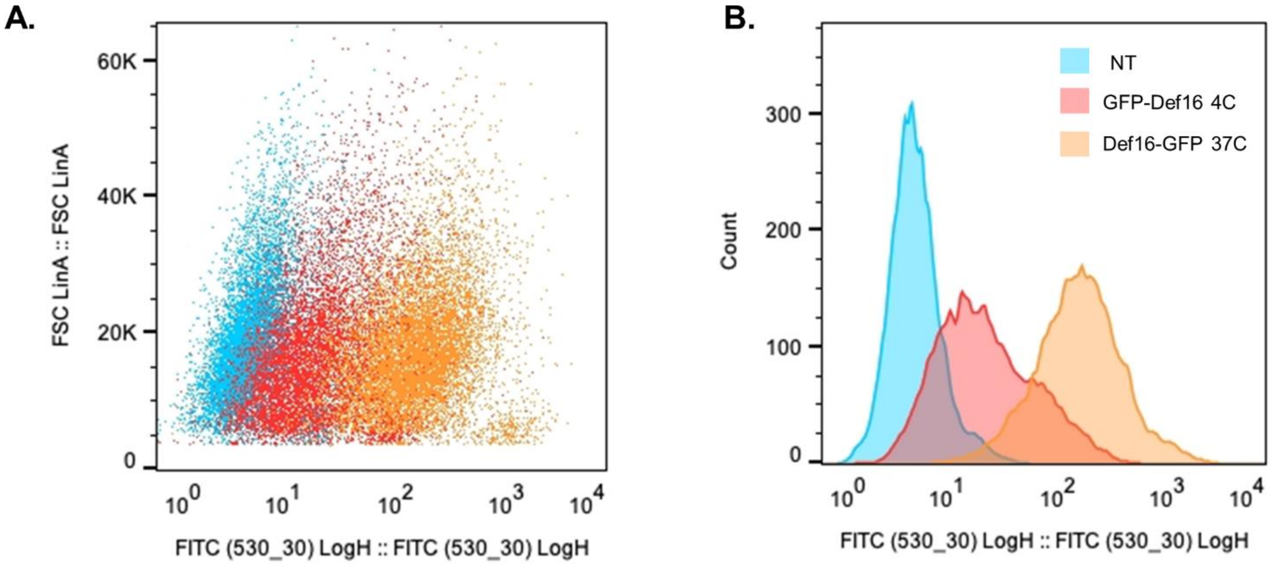


**Figure S5: Flow Cytometry Analysis Def16-GFP and GFP-Def16 in HeLa Cells.** HeLa cells were incubated with recombinant proteins at a concentration of 5 μM for two hours at 37 ℃ and 4 ℃. Subsequently, the cells were washed with PBS and trypsinized. The cells were then resuspended in PBS supplemented with 2.5% FBS. **(A)** The Y-axis represents forward scatter size (FSC), where an increased signal indicates an increase in cell size or budding. The X-axis represents the GFP signal in different cell populations. **(B)** The histogram displays the count of cells versus GFP intensity for each treatment.

**Supplementary Figure S6**

To quantify the extent of Def16-GFP penetration into cells, HeLa cells were incubated with variable concentrations of the protein for two hours at 37°C. Subsequently, the cells were washed with PBS, fixed, and stained with DAPI (blue) and β-tubulin (red) for visualization using confocal imaging (Fig. S6-A). To assess the percentage of cellular permeability, we utilized a threshold module in Fiji software to quantify the green and red channels. The selected green areas were then divided by the total cellular surface area indicated by the red channel (β-tubulin). These values were normalized relative to the mean green/red fluorescence intensity ratio of untreated cells, revealing a dose-dependent signal for Def16-GFP penetration into the cells (Fig. S6-B). Furthermore, we calculated the percentage of cells penetrated by Def16-GFP by “Coloc 2” plugin (Fiji), and determined the Pearson's coefficient, which provides a statistical measure of the linear relationship between the green and red channels at different Def16-GFP concentrations (Fig. S6-C). Notably, a clear correlation is observed, indicating a penetration rate of nearly 100% in cells treated with 10 μM of the protein.

**A.**

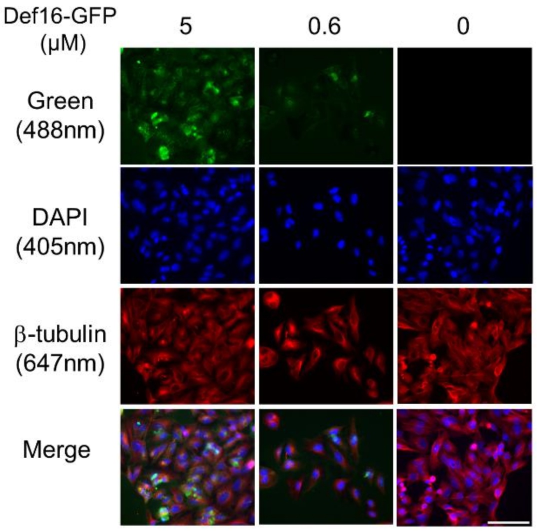


**B.**

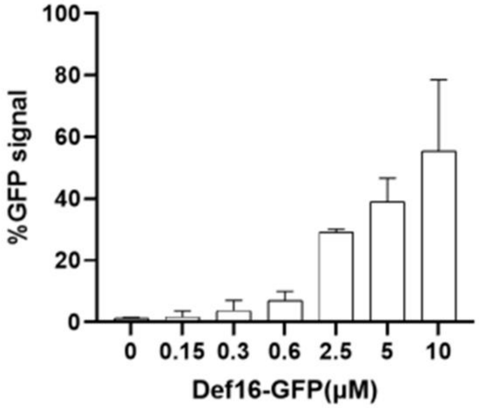


**C.**

| Def16-GFP (μM) | % Cellular uptake | Pearson coefficient |
|---|---|---|
| 10 | 99.3 | 0.52±0.14 |
| 5 | 90.7 | 0.39 |
| 2.5 | 19.02 | 0.35 |
| 0.3 | 0 | 0.335±0.06 |
| 0 | 0 | 0.095±0.007 |

**Figure S6. Cellular Delivery of Def-GFP in HeLa Cells. (A)** HeLa cells were incubated with variable concentrations of Def16-GFP for 2 hours. Cells were subsequently stained and imaged for DAPI (blue), Def16-GFP (green), and β-tubulin (red), and the cellular uptake of the protein was analyzed. Representative fields were imaged at 40× magnification. Scale bar: 100 µm. **(B)** Quantification of confocal image signal intensity. The bars represent the mean green fluorescence intensity relative to non-treated cells, normalized to the cellular surface area measured by β-tubulin. Values were measured using the Color Threshold tool in Fiji. The values represent the mean ± SD of four replicates. **(C)** Measurement of protein penetration levels. The table summarizes Pearson's coefficient and the cellular uptake percentage of Def16-GFP. The latter was calculated based on the number of green pixels that colocalized with the cellular margins of red pixels (β-tubulin) and were above the threshold in both channels. The Pearson's coefficient for channel-1 (green) vs. channel-2 (red) was measured using the "Coloc 2" plugin in Fiji and is presented as the mean ± SD of duplicates.
